# TRAF6 coiled-coil domain mediates its trimerization

**DOI:** 10.64898/2025.12.24.696318

**Authors:** Mohammed Khan, Rui Yang, Qian Yin

**Affiliations:** Department of Biological Science, Florida State University, Tallahassee, FL 32306; Institute of Molecular Biophysics, Florida State University, Tallahassee, FL 32306

**Keywords:** coiled-coil domain, oligomerization, TRAF6, trimerization

## Abstract

Tumor necrosis factor receptor-associated factor 6 (TRAF6) is an adaptor protein that plays a critical role in innate immune signaling. TRAF6 consists of an N-terminal RING domain, five zinc fingers, a coiled-coil domain, and a C-terminal TRAF domain. Although structural and oligomeric information for coiled-coiled domains in TRAF2-5 is known, information on TRAF6’s coiled-coiled domain is relatively unknown. Here, we characterized the oligomeric state of the human TRAF6 domain (residues 288-348) using size-exclusion chromatography (SEC) and size-exclusion chromatography coupled multi-angle light scattering (SEC-MALS). Both His-tagged and tagless proteins displayed molecular masses consistent with a trimer. Coiled-coil computational tools identified a heptad repeat pattern that supported the trimeric findings. AlphaFold-Multimer modelling further showed a symmetric, trimeric coiled-coiled domain with a conserved hydrophobic core. These results provide experimental support for TRAF6 coiled-coiled domain trimerization and help establish a structural framework for understanding how TRAF6 assembles into higher-order structures.

## Introduction

Tumor necrosis factor (TNF) receptor-associated factors (TRAFs) are intracellular signaling proteins that transduce signals from cell-surface receptors, including TNF receptors (TNFRs), Toll-like receptors (TLRs), and interleukin receptors (1). Except for TRAF1, which does not contain a RING domain, all TRAF proteins are composed of an N-terminal RING finger followed by a number of zinc fingers (ZnFs), a central coiled-coil domain (CC), and a C-terminal TRAF domain (Figure 1A). The coiled-coil and the TRAF domain are also known as TRAF-N and TRAF-C domains (2).

**Figure 1.**
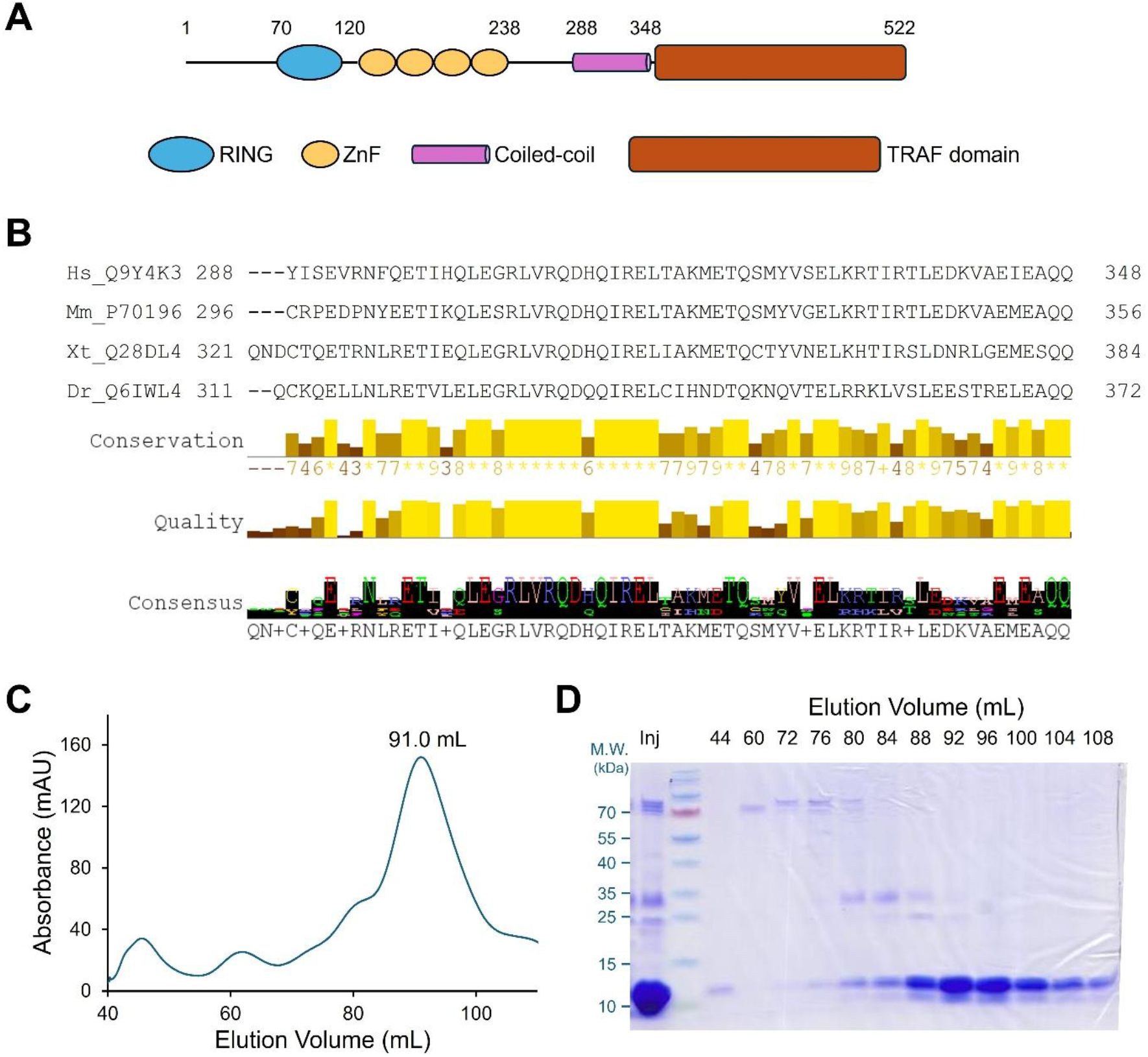
Purification of TRAF6 coiled-coil region (TRAF6-CC). (**A**) A schematic diagram of human TRAF6 protein. The domain boundaries are marked on top of the diagram. ZnF: zinc finger; (**B**) Sequence alignment of human (Hs), mouse (Mm), frog (Xt), and drosophila (Dr) TRAF6 coiled-coil regions. The UniProt identifiers and residue numbers are marked. Conservation and consensus residues are generated by JalView (34); (**C**) Size-exclusion chromatography profile of His6-TRAF6-CC from a HiLoad 16/600 Superdex 200 column; (**D**) SDS-PAGE analysis of His6-TRAF6-CC from the SEC profile in (**C**). Molecular markers are labeled to the left of the gel. Inj: SEC injection/input sample.

RING domains often function as E3 ubiquitin ligases, adding polyubiquitin chains to a variety of substrates (3). An increasing number of RING domains are shown to homo- or hetero-dimerize to mediate ubiquitin chain transfer (4, 5). Although all RING-containing TRAF proteins were presumed to possess E3 ligase activity, only TRAF6 has been experimentally validated to catalyze polyubiquitination (5, 6). Together with the heterodimeric E2 ubiquitin-conjugating enzyme Ubc13/Uev1A, TRAF6 adds lysine 63 (K63)-mediated polyubiquitin chain to itself and target proteins, creating a platform for recruitment of downstream signaling molecules (7). The first zinc finger of TRAF6, though not directly interacting with the E2, substantially enhances TRAF6 E3 ligase activity by positioning the ubiquitin in the E2∼ubiquitin conjugate (6). In contrast, the highly homologous TRAF2 does not interact with Ubc13 or mediate polyubiquitination because of amino acid alterations in the RING domain (8).

In TRAF2, 3, 4, and 5, the coiled-coil domain mediates TRAF trimerization together with the C-terminal TRAF domain (9-15). Interestingly, while the C-terminal TRAF domain trimerizes, the N-terminal RING domain forms dimers (5, 6, 8). This seemingly stoichiometry incompatibility, however, is proposed to drive the formation of a higher-order hexagonal honeycomb pattern that facilitates the recruitment of downstream proteins and enhances signaling (5, 6, 16).

Oligomerization and higher-order interactions are central to TRAF signaling, making the structure and mechanism of the coiled-coil domain critical to TRAF function. Coiled-coil domains are alpha-helical structures that oligomerize through intermolecular interactions. Depending on the amino acid composition, coiled-coil domains may dimerize, trimerize, or tetramerize (17, 18). Coiled-coils can adopt a variety of configurations, including homo-oligomers or hetero-oligomers, parallel or anti-parallel, and obligated or dissociative (17, 18). The versatility and dynamic nature of coiled-coil organization can be found in many multi-domain proteins and multi-subunit protein complexes, mediating crucial cellular functions such as transcription, cell motility, membrane fusion, and protein quality control (17-19). The organizational and structural building block in naturally occurring, left-handed coiled-coils is the heptad, *abcdefg*. Positions *a* and *d* form the hydrophobic core, and they are usually occupied by leucines, isoleucines, and to a lesser degree, valines. The other positions contribute to coiled-coil stabilization and interaction with the environment, and they can be occupied by both hydrophobic and hydrophilic residues (17, 18). The continuity of coiled-coils may be disrupted by noncanonical residues, leading to changes in periodicity and flexibility in coiled-coils.

Structural information is available for the RING and zinc fingers of TRAF2, TRAF5, and TRAF6, both alone and in complex with E2 and E2∼Ub conjugate (5, 6, 8, 20). The TRAF domains of TRAF2, 3, 4, 5, and 6 in complex with various ligand peptides or proteins have been extensively characterized (9-15, 21-24). Multiple TRAF2, 3, 4, and 5 TRAF domain structures encompass the entire or partial coiled-coil domain (9-15, 23), revealing coiled-coil domain-mediated trimerization. Crystal structures of TRAF2 coiled-coil homotrimer and TRAF1:TRAF2 coiled-coil heterotrimer are also reported (25). Despite extensive structural characterization of TRAF family domains, the oligomeric organization and high-resolution structural information on the TRAF6 coiled-coil domain remain unresolved. In the absence of the coiled-coil domain, the TRAF6 TRAF domain is monomeric (21), highlighting the essential role of the coiled-coil domain in mediating TRAF6 oligomerization. Here, we biochemically characterized human TRAF6 coiled-coil domain (TRAF6-CC) by size-exclusion chromatography (SEC) and size-exclusion chromatography coupled multi-angle light scattering (SEC-MALS), confirming the trimeric organization of TRAF6-CC. We further analyzed TRAF6-CC via coiled-coil domain prediction tools, identifying heptad patterns and key interfacial residues. Lastly, we modeled the trimeric structure of TRAF6-CC using AlphaFold Multimer, completing a missing piece of the TRAF6 structure.

## Results

### TRAF6 coiled-coil domain forms a trimer in solution

Human TRAF6 is a 522-amino acid protein composed of an N-terminal RING domain (70-120), four zinc fingers (121-238), a coiled-coil domain (288-348), and a C-terminal TRAF domain (349-504) (Figure 1A). The RING and the first zinc finger are crucial for the E3 ubiquitin ligase activity (5, 6), while the TRAF domain interacts with receptors and adaptor proteins (21). The coiled-coil domain of TRAF6 is highly conserved across the vertebrate homologs, including both hydrophobic and hydrophilic residues (Figure 1B). For biochemical characterization, we expressed and purified TRAF6-CC (residues 288-348) from *E. coli* using affinity purification followed by size-exclusion chromatography (SEC). The SEC profile displayed a sharp, symmetrical peak (Figure 1C), indicating a well-folded, homogeneous protein. SDS-PAGE analysis of SEC fractions showed a major band of ∼ 10 kDa in the SEC peak fractions (Figure 1D), further confirming the identity, purity, and homogeneity of the purified TRAF6-CC.

The molecular mass of TRAF6-CC with the N-terminal His6 tag is 9.61 kDa. However, it eluted from the HiLoad 16/600 SEC column at the volume of 91.2 mL (Figure 1C), corresponding to a molecular mass of ∼ 20 kDa based on column calibration. The elution volume suggests an oligomeric organization of TRAF6-CC. To further investigate the oligomeric status of TRAF6-CC, we performed size-exclusion chromatography coupled multi-angle light scattering (SEC-MALS), a shape-independent technique that measures the molecular mass of protein assemblies in solution. We performed SEC-MALS on both hexahistidine-tagged TRAF6-CC (His_6_-TRAF6-CC) and tagless TRAF6-CC (Figure 2A-B). The SEC-MALS experiments revealed a molecular mass of 25.6 kDa (±3.6%), consistent with a trimeric molecular mass of 28.8 kDa (Figure 2A-B). Similarly, the tagless TRAF6-CC measured 20.9 kDa from SEC-MALS, consistent with a trimeric molecular mass of 23.2 kDa (±2.8%) (Figure 2B).

**Figure 2.**
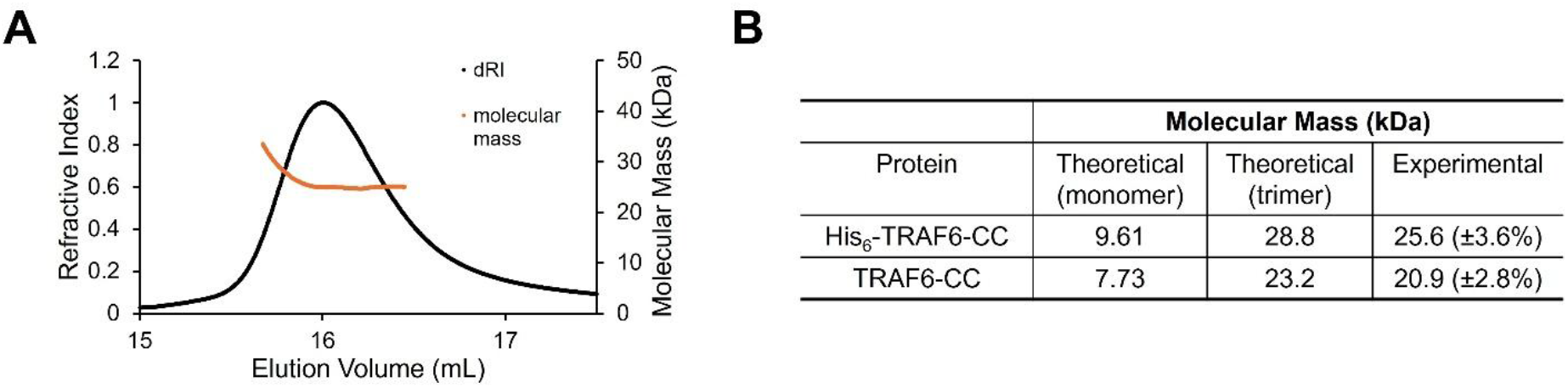
SEC-MALS determination of TRAF6-CC molecular mass. (**A**) A representative SEC-MALS profile of His_6_-TRAF6-CC. Black and orange traces represent the refractive index of His_6_-TRAF6-CC peak and calculated molecular mass across the peak, respectively. (**B**) Summary of theoretical and experimentally determined molecular mass of His_6_-TRAF6-CC and tagless TRAF6-CC. In the parentheses are the data fitting errors.

### Modeling TRAF6 coiled-coil trimer organization

The SEC-MALS measurement establishes that TRAF6 coiled-coil domain (TRAF6-CC) adopts a trimeric organization in solution, the same way TRAF2, 3, 4, and 5 coiled-coil domains oligomerize (9-15, 25). In the structures, the coiled-coil domains of TRAF2-5 form a typical parallel coiled-coil mediated by hydrophobic residues at positions *a* and *d* of the heptad repeats. As no high-resolution structural information is available for TRAF6-CC, we employed a few computational tools to predict the alpha-helical content and coiled-coil probability of TRAF6-CC. Both MarCoil and MultiCoil2 (26, 27) predict that the TRAF6-CC region is completely alpha-helical. The two algorithms also predict that the majority of TRAF6-CC forms continuous coiled-coil without interruption or perturbation, with the region encompassing N295-E345 scoring >90% probability (Figure 3A-B). The agreement between the two algorithms reinforces the likelihood of coiled-coil formation. Each TRAF6-CC molecule is predicted to fold into an amphipathic alpha helix (Figure 3C-D). MultiCoil2 also predicts that TRAF6-CC has a higher probability of forming a trimeric coiled-coil (Figure 3D), consistent with our SEC-MALS results. TRAF6-CC region is predicted to contain seven complete heptads, starting from I299 and ending at Q347. I299, L306, I313, M320, V327, I334, and V341 occupy the *a* positions; while L302, Q309, L316, Q323, L330, L337, and I344 occupy the *d* positions (Figure 3C-D). These hydrophobic residues constitute the core of the trimer coiled-coil interface. MarCoil also predicted that the charged residues, including D310, R314, E315, K319, E329, R332, R335, E338, D339, K340, E343, and E345, form intrahelical interactions to stabilize the alpha helices (Figure 3C).

**Figure 3.**
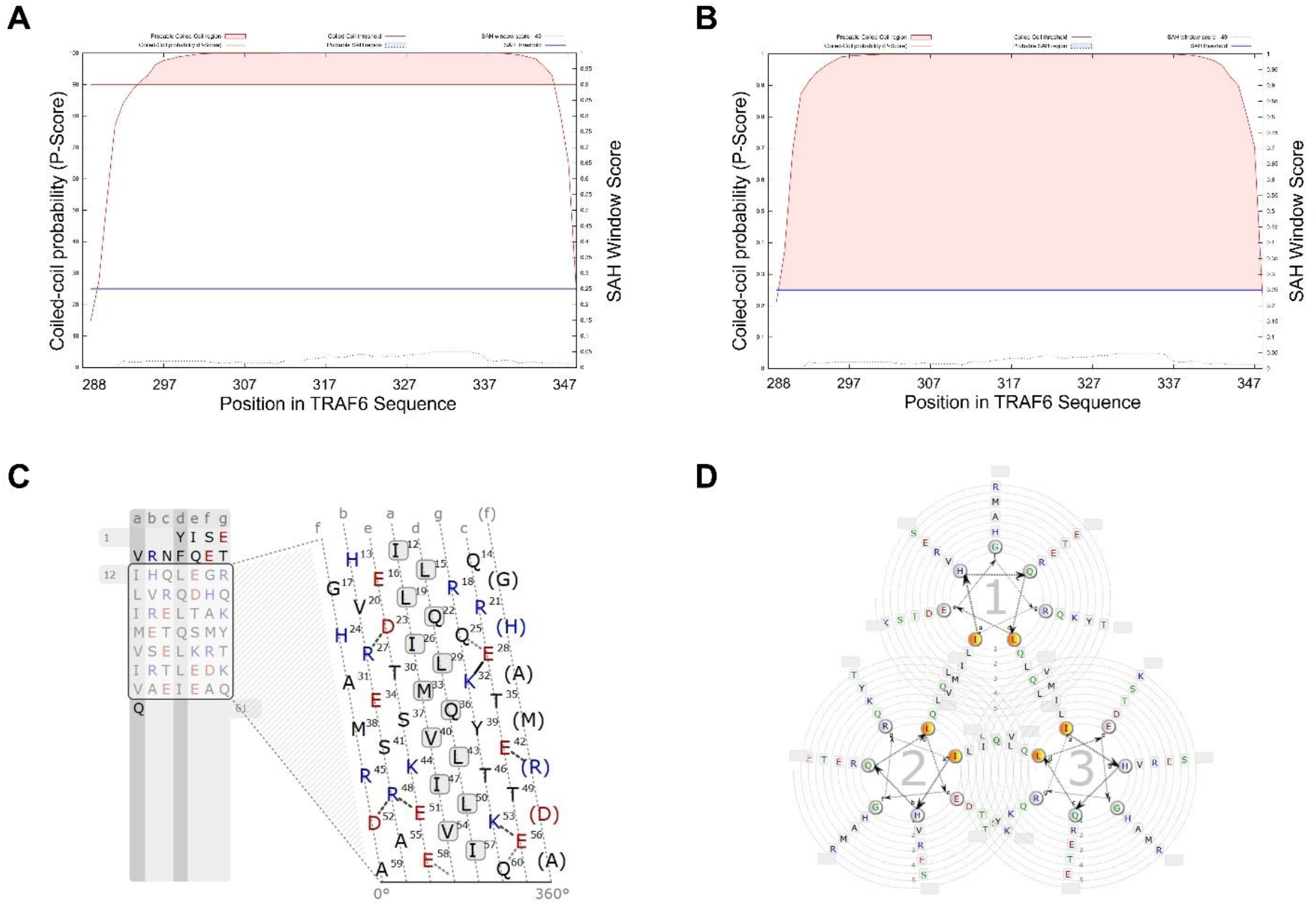
Coiled-coil prediction of TRAF6-CC. (**A**) and (**B**) Coiled-coil probability and single alpha helix (SAH) content predicted by MarCoil (**A**) and MultiCoil2 (**B**). Red shaded area represents predicted coiled-coil region along TRAF6-CC sequence; (**C**) Helical projection of TRAF6-CC. Hydrophobic and polar residues are colored black, positively charged residues are colored blue, and negatively charged residues are colored red. The black box marks predicted coiled-coil sequence, and residues are arranged in repeated heptads. Predicted electrostatic interactions are marked by solid and dashed lines. The residues in (**C**) are numbered based on their positions in the coiled-coil region; to convert to positions in full-length TRAF6, add 287 to the number; (**D**) Helical wheel representation of the TRAF6-CC trimer. The figure is generated by the waggawagga webserver (32).

To further understand how the residues pack interact with each other to form the trimer, we modeled the TRAF6-CC trimer using the AlphaFold2 server (28, 29). The predicted trimer model revealed a symmetric, parallel, triple-stranded coiled-coil spanning ∼ 85 Å (Figure 4A). Each TRAF6-CC is a continuous, curved alpha helix. The interfacial residues predicted by MarCoil and Multicoil2 point inward, forming the trimer’s hydrophobic core (Figure 4B). The trimer model also revealed potential interhelical interactions that are not present in the coiled-coil predictions. For example, D310, instead of interacting with R314 in the same helix, forms a salt bridge with R305 on a neighboring helix, likely bolstering the stability of the coiled coil (Figure 4C). This discrepancy likely arises from the curved nature of the alpha helices in the trimeric coiled-coil. Most of the interfacial residues are conserved across species (Figure 1B), underscoring their importance in TRAF6 function, likely through mediating trimerization.

**Figure 4.**
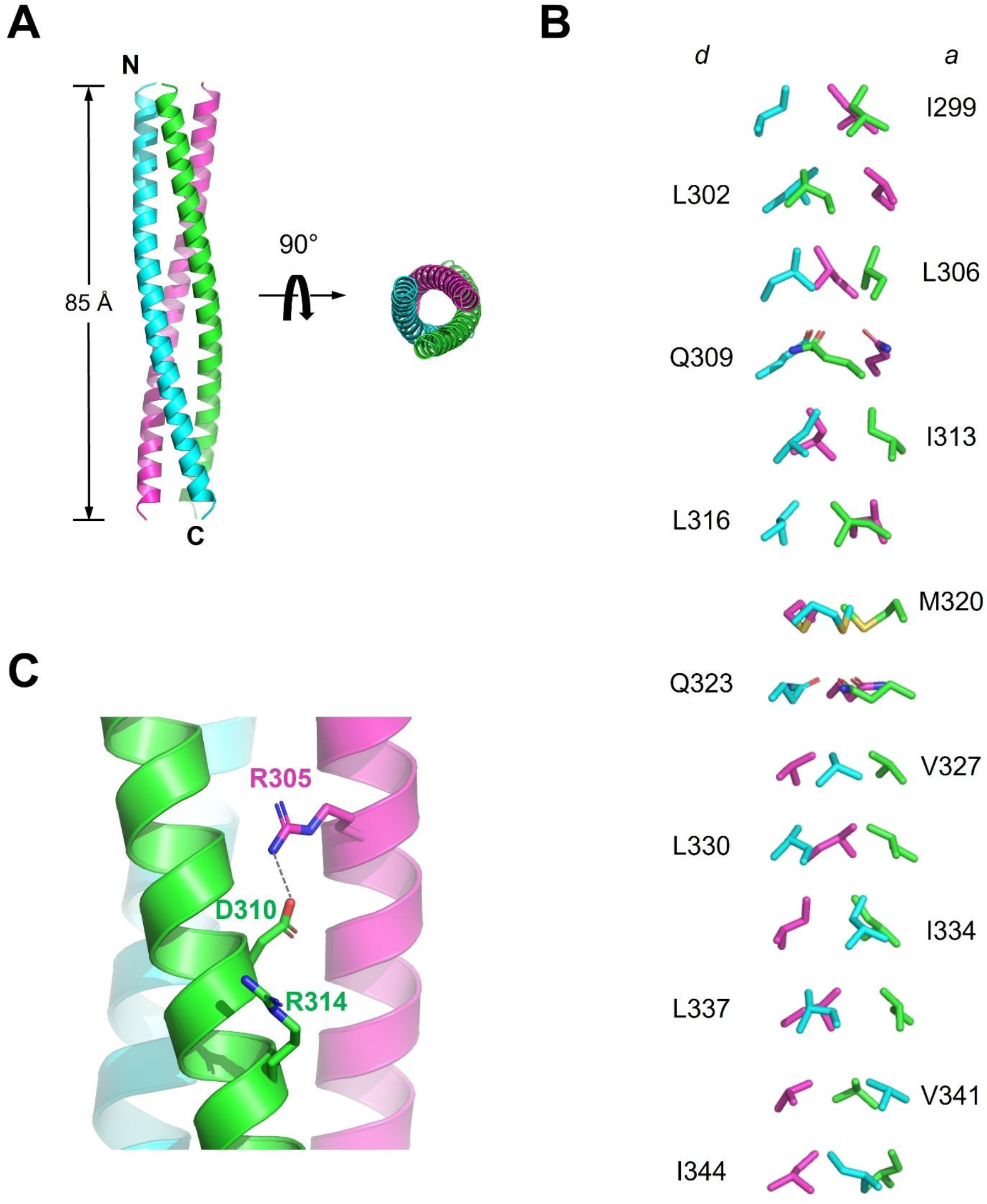
Structure model of TRAF6-CC trimer. (**A**) Cartoon representation of AlphaFold generated TRAF6-CC trimer in two orthogonal orientations. The three strands are colored green, cyan, and magenta, respectively. The N- and C-termini of the cyan strand are marked; (**B**) A detailed view of the *a* and *d* position residues. For clarity, the rest of the coiled-coil is omitted; (**C**) A zoom-in view of the predicted salt bridge between D310 of one strand (green) and R305 of a neighboring strand (magenta).

## Discussion

In this work, we experimentally established the oligomerization state of the predicted TRAF6 coiled-coil region, confirming its trimeric organization, like other human TRAF proteins (9-15, 25). Computational modelling supports a trimeric, parallel coiled-coiled structure that aligns with our SEC-MALS results. The hydrophobic residues predicted to form the trimer core are highly conserved across species, underscoring their importance in TRAF6 function, likely through mediating trimerization.

Both the coiled-coil prediction algorithms and AlphaFold Multimer placed certain noncanonical residues at the *a* and *d* positions, including Q309, M320, and Q323. These residues contain side chains bulkier than canonical leucines and isoleucines. To accommodate these residues, the coiled-coil is expected to increase in diameter at such positions. While the AlphaFold model prediction is likely to largely represent the true structure, it does not fully take into account the noncanonical residues present at the trimer’s interface. Indeed, in the current model, the three M320 residues clash into each other. Therefore, an experimentally determined TRAF6-CC structure is still of importance to fully reveal its trimeric organization in atomic detail.

The homotrimeric TRAF2 coiled-coil domain and the heterotrimeric TRAF1:TRAF2 coiled-coil interact with the BIR1 domain of the protein cell inhibitor of apoptosis 2 (cIAP2) [ref]. However, all three molecules in the TRAF2 coiled-coil trimer differ in their conformation, an asymmetry likely introduced by interaction with cIAP2. TRAF6-CC has been reported to interact with Ubc13 (30, 31). It will be interesting to investigate the stoichiometry of this interaction and potential conformational changes in TRAF6-CC.

## Materials and Methods

### Protein Expression and Purification

The region encoding the human TRAF6 (UniProt Q9Y4K3) coiled-coil domain (Y288-Q348) was cloned into a pET28a vector (Novagen) between the *Nde*I and *Xho*I sites using standard PCR protocols. The fusion protein contains an N-terminus His_6_ tag followed by a thrombin cleavage site and the TRAF6 coiled-coil domain (His_6_-THROM-TRAF6-CC, abbreviated as His_6_-TRAF6-CC). This sequence was validated by Sanger sequencing. The expression plasmid was transformed into *E. coli* BL21-CodonPlus (DE3) RIPL cells (Agilent) and grown at 37°C. The bacterial culture was induced with 0.4 mM IPTG when the optical density (OD600) reached ∼0.7, and the cells were incubated at 18°C overnight. The cells were harvested by centrifugation at 4000 *g* at 4°C for 20 min. The cell pellet was resuspended in Lysis Buffer (50 mM sodium phosphate pH 7.4, 300 mM NaCl, 20 mM imidazole, 5 mM β-mercaptoethanol, and 10% glycerol). Cells were lysed by sonication, and the cell debris was removed by centrifugation at 18,000 *g* for 60 min at 4°C. His_6_-TRAF6-CC was purified from the supernatant by affinity chromatography using Ni-NTA resin (Qiagen). The resin was washed with 10 column volumes of Wash Buffer (50 mM sodium phosphate pH 7.4, 300 mM NaCl, 30 mM imidazole and 5 mM β-mercaptoethanol) and the bound proteins were eluted using Elution Buffer (50 mM sodium phosphate pH 6.0, 300 mM NaCl, 300 mM imidazole, and 5 mM β-mercaptoethanol). Fractions containing His_6_-TRAF6-CC were pooled and loaded onto a Superdex 200 HiLoad 16/600 size-exclusion column (GE Life Sciences) to separate the protein from other impurities. The column was pre-equilibrated with Running Buffer (20 mM Tris-HCl pH 8.0, 150 mM NaCl, and 5 mM β-mercaptoethanol). Peak fractions were collected, pooled, and concentrated to 9.83 mg/ml. All purified proteins were immediately frozen in liquid nitrogen and stored at −80°C. The N-terminal His_6_ tag was removed by incubating His_6_-TRAF-CC with bovine thrombin protease (BioVision) at 20°C for overnight. The cleaved His6 tag was removed by reverse Ni-NTA affinity chromatography. The flow-through fraction containing tagless TRAF6-CC was further purified by SEC as described above.

### Size-Exclusion Chromatography Coupled Multi-Angle Light Scattering (SEC-MALS)

For SEC-MALS, TRAF6-CC was analysed with and without its His_6_ tag (His_6_-TRAF6-CC and TRAF6-CC, respectively). The concentrations of His_6_-TRAF6-CC were 1 mg/ml and 6.57 mg/ml. The TRAF6-CC concentrations were 0.934 mg/ml and 1.57 mg/ml. The SEC-MALS experiment was calibrated using bovine serum albumin (BSA) (Millipore Sigma). Protein samples were injected into a Superdex 200 Increase 10/300 size-exclusion column equilibrated in a buffer containing 20 mM Tris-HCl pH 8.0 and 150 mM NaCl. The column was coupled to a multi-angle light scattering detector (DAWN HELEOS II, Wyatt Technology) and a refractometer (Optilab T-rEX refractometer, Wyatt Technology). The flow rate was 0.5 mL/min. Data processing was carried out using the program ASTRA 7.3 (Wyatt Technology). Molar mass and mass distribution of each protein were calculated and reported by ASTRA 7.3.

### Coiled-coil analysis

The coiled-coil analysis was performed using the Waggawagga webserver (32). The entire TRAF6-CC sequence Y288-Q348 was uploaded to the Waggawagga server. Coiled-coil Tools include Marcoil and Multicoil2, Ncoils, and Paircoils. We set the window length as 21.

### Structural modeling

The trimeric model of TRAF6-CC was generated by AlphaFold Multimer (29) using the Cosmic 2 interface and default parameters. The trimer model was visualized using PyMol (33).

## Acknowledgments

We thank Dr. Peter Randolph at the Physical Biochemistry Facility at FSU for assistance in SEC-MALS analysis. We thank Dr. Gwimoon Seo at the Protein Expression Facility at FSU for competent cells. We thank Dr. Madhurima Bhattacharya for assistance in protein expression and purification. This work is supported by NIH grant R01AI146330 (Q.Y.).

